# Tissue of origin detection for cancer tumor using low-depth cfDNA samples through combination of tumor-specific methylation atlas and genome-wide methylation density in graph convolutional neural networks

**DOI:** 10.1101/2023.12.24.573277

**Authors:** Trong Hieu Nguyen, Nhu Nhat Tan Doan, Trung Hieu Tran, Le Anh Khoa Huynh, Phuoc Loc Doan, Thi Hue Hanh Nguyen, Van Thien Chi Nguyen, Giang Thi Huong Nguyen, Hoai-Nghia Nguyen, Hoa Giang, Le Son Tran, Minh Duy Phan

## Abstract

**Background:** Cell free DNA (cfDNA)-based assays hold great potential in detecting early cancer signals yet determining the tissue-of-origin (TOO) for cancer signals remains a challenging task. Here, we investigated the contribution of a methylation atlas to TOO detection in low depth cfDNA samples.

**Methods:** We constructed a tumor-specific methylation atlas (TSMA) using whole-genome bisulfite sequencing (WGBS) data from five types of tumor tissues (breast, colorectal, gastric, liver and lung cancer) and paired white blood cells (WBC). TSMA was used with a non-negative least square matrix factorization (NNLS) deconvolution algorithm to identify the abundance of tumor tissue types in a WGBS sample. We showed that TSMA worked well with tumor tissue but struggled with cfDNA samples due to the overwhelming amount of WBC-derived DNA. To construct a model for TOO, we adopted the multi-modal strategy and used as inputs the combination of deconvolution scores from TSMA with other features of cfDNA.

**Results:** Our final model comprised of a graph convolutional neural network using deconvolution scores and genome-wide methylation density features, which achieved an accuracy of 69% in a held-out validation dataset of 239 low-depth cfDNA samples.

**Conclusions:** In conclusion, we have demonstrated that our TSMA in combination with other cfDNA features can improve TOO detection in low-depth cfDNA samples.

## Background

Liquid biopsies based on cell free DNA (cfDNA) have recently emerged as a novel method for early cancer detection owing to their non-invasive, sensitive, and multi-modal characteristics. Multiple features can be derived from cfDNA sequences to reveal various aspects of cancer-specific aberration including fragment length profile [1]–[3], copy number aberration [4], motif- end [5], genome-wide and targeted methylation profiles [6]. To maximize the capacity to distinguish between cancer patients and healthy individuals, the integration of multiple features in advanced machine learning or deep learning has become a common approach and demonstrated encouraging performance [7]–[12]. However, predicting tumor of origin (TOO) for multiple cancer types at early stage remains challenging due to the low abundance of cfDNA originating from tumors (ctDNA), further confounded by the presence of various DNA components released from non-tumor sources.

Recent advances have highlighted the increasing importance of DNA methylation in the early cancer detection context [13]–[15]. DNA methylation is an epigenetic marker that plays a crucial role in regulating gene expression, maintaining genomic stability, and directing cell development. These epigenetic modifications contribute heavily to the dysregulation of multiple pathways, allowing cancer cells to proliferate uncontrollably [16]–[19]. Most importantly, these DNA methylation patterns are tissue-specific and remain stable during neoplastic transformation, which could allow tumor of origin identification [20]–[22]. Therefore, characterization of DNA methylation biomarkers in tumor tissues could potentially guide the detection of both cancer characteristics and TOO in cfDNA samples.

Constructing a methylation atlas is an attractive approach in analyzing large methylation data. Moss *et al*. [23] developed a comprehensive methylation atlas of healthy human cell types using data from 450K Illumina microarray to decompose of human cell types in bulk samples. Loyer *et al*. [24] created a human methylation atlas using deep whole-genome bisulfite sequencing (WGBS) from 39 normal cell types of 205 healthy tissue samples to further enhance the understanding of the human normal cell types methylome. The methylation atlas was constructed based on differential fractions of hypo-methylated and hyper-methylated reads at various cell-type specific genome segments. While the authors reported a promising deconvolution resolution of 0.1%, the study did not demonstrate the application of this methylation atlas to determine TOO of cfDNA samples from cancer patients.

In this study, we investigated the application of methylation atlas to cell type deconvolution of cfDNA samples, with the ultimate goal of creating a model that can detect TOO in low-depth cfDNA samples for early cancer screening (Figure 1). We first constructed a tumor-specific methylation atlas (TSMA) using WGBS data from five types of tumor tissues (breast, colorectal, gastric, liver and lung cancer) and paired white blood cells (WBC). We then validated the use of TSMA in deconvolution of tumor tissue samples and cfDNA samples, demonstrated that TSMA worked well for tumor tissues but struggled with cfDNA samples due to the overwhelming amount of DNA fragments derived from WBC. To construct a model for TOO in low-depth cfDNA samples, we adopted the multi-modal strategy and combined deconvolution scores from our TSMA with other features previously explored by our group [9]. Our final model comprised of a graph convolutional neural network (GCNN) using deconvolution scores and genome-wide methylation density features (GWMD) as inputs and achieved an accuracy of 69% in a held-out validation dataset of 239 samples.

**Figure 1.**
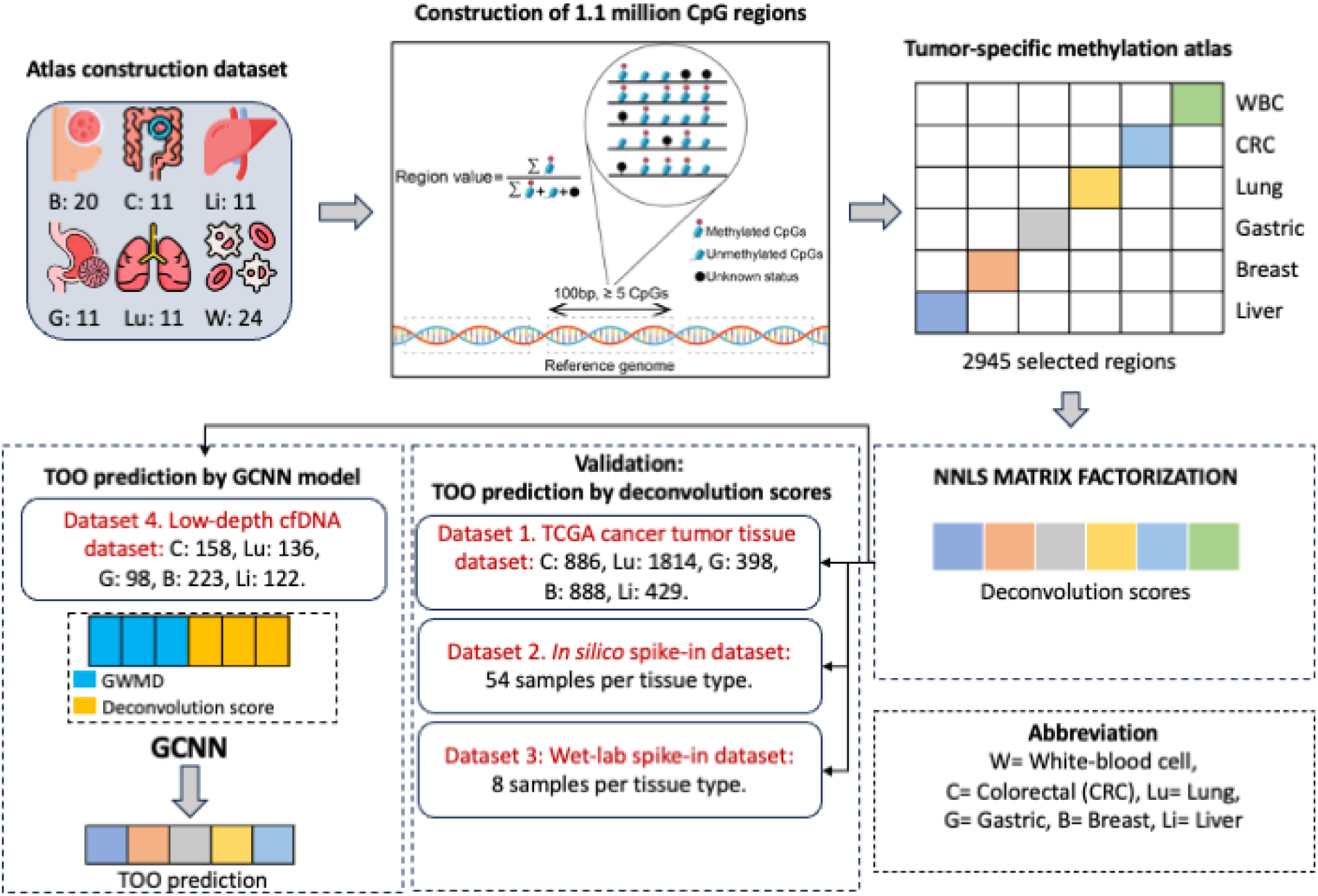
Schematic overview of this study. We first constructed a tumor-specific methylation atlas (TSMA) using a WGBS dataset (Atlas construction dataset) of 64 tumor tissues comprising five cancer types (breast, colorectal, gastric, liver and lung cancer) and paired white blood cells (WBC). The methylation signals from WGBS data (region value) were calculated for approximately 1.1 million pre-defined CpG regions (regions of 100bp in length covering at least 5 CpG sites). This large matrix of region values was then used to construct the TSMA, comprising of 2,945 differential regions between five tumor-tissue types and WBC across the entire genome. With the TSMA, deconvolution scores for new input samples were calculated using a non-negative least square (NNLS) matrix factorization. We next validated the use of TSMA and deconvolution scores in 3 datasets, including Dataset 1: tumor tissue methylation microarray data from TCGA, Dataset 2: in silico spike-in samples with known amount of tissue DNA fragments and Dataset 3: wet lab spike-in samples with known amount of tissue DNA fragments. Finally, we implemented a graph convolutional neural network combining deconvolution scores and genome-wide methylation density (GWMD). The model was trained and validated on a cohort of 737 low-depth WGBS cfDNA samples (Dataset 4).

## Materials and methods

### Dataset description

#### Atlas construction dataset

This dataset includes 64 tumor tissue samples from five distinct cancer classes (Liver (11 samples), Breast (20 samples), Lung (11 samples), CRC (11 samples), Gastric (11 samples)) and 24 WBC samples. All samples were whole-genome bisulfite sequenced at 5-15x depth coverage. Metadata is provided in Supplementary Table 2.

Validations were conducted on four different datasets:

- ***Dataset 1:*** 450K/850 methylation microarray datasets were downloaded from TCGA. The curated dataset only includes samples from our five group of tissues of interest (4,415 samples), which consists of 886 CRC samples, 1,814 Lung samples, 398 Gastric samples, 888 Breast samples and 429 Liver samples from cancer patients.
- ***Dataset 2:*** We generated an *in-silico* spike-in dataset using three healthy WGBS cfDNA samples as background. To simulate the presence of circulating cell-free tumor-derived DNA (ctDNA) at varying abundance levels, spike-in reads were randomly extracted from pooled tumor tissue samples in the Atlas dataset, representing ratios of 0.01%, 0.05%, 0.1%, 1%, 10%, and 25%. These simulated reads were then merged with the background cfDNA samples, generating 3 samples for each cancer type at each spike-in ratio (3 samples x 5 cancer types x 6 ratios = 90 samples). The process was repeated three times, generating in total a dataset of 270 samples. This simulation dataset allows us to assess the sensitivity of our constructed atlas in detecting tumor-related signals within the cfDNA background at different levels of abundance.
- ***Dataset 3:*** We refer to section “Wet-lab spike-in experiments in Dataset 3” in Materials and methods for a detailed description of this dataset.
- ***Dataset 4***: A cohort of 737 low-depth WGBS cfDNA samples (0.5x). The low-depth dataset was previously employed in the construction and validation of an integrated multi-modal model for early cancer detection [9]. 498 samples are used in the training set and 239 samples are served as a held-out validation set.

Metadata tables for all datasets are provided in Supplementary Table 3.

### Wet-lab spike-in experiments in Dataset 3

This section is devoted to the preparation of Dataset 3. The spike-in experiment was conducted using healthy control and cancerous tissue samples. The healthy control sample was created by pooling cfDNA of multiple healthy, non-cancerous individuals. The extracted cancer gDNA was subjected to a fragmentation process to create fragmented cancer DNA. These cancer DNA fragments were then spiked in the healthy control sample with four different amounts to create a set of four samples containing different tumor abundances (0.1%, 1%, 10% and 25% of cancer DNA in total amount of DNA). This experiment was repeated twice for each cancer type with extracted gDNA from two different cancer samples, resulting in a total of 40 samples (10 cancer samples in total – two samples for each type of cancer including breast, colorectal, liver, lung and gastric cancer). These spike-in samples then undergone bisulfite conversion and purification by EZ DNA Methylation-Gold Kit (Zymo Research, D5006, USA). DNA library was prepared from bisulfite-converted DNA samples using xGen Methyl-Seq DNA Library Prep Kit (Integrated DNA Technologies, 10009824, USA) with Adaptase technology, according to the manufacturer’s instructions. All the DNA concentrations were identified by the QuantiFluor dsDNA system (Promega, USA). The library products were sequenced on the DNBSEQ-T7 DNA system (MGI Tech, Shenzhen, China).

### Bioinformatics pipeline

FASTQ files were fed to an in-house Bioinformatics pipeline. We performed FASTQ file quality control with fastqc (version 0.11.2 [34]) and trimming adapters and low-quality bases by TrimGalore (version 0.6.7, [35]). We only trimmed the first 15bp --HEADCROP at the 5’ end of Read 1, since it is not possible to exactly locate the 5’ end of read 2 in this assay. Sequence alignment and CpG site methylation calling were done by the Bismark suite (version 0.23.1, [36]). We used picard MarkDuplicate (version 2.18.7 [37]) to mark and remove duplicated reads. Reads were then filtered by samtools (version 1.18, [38]) to keep only reads whose quality is greater than Q30. All tools’ parameters were kept default unless mentioned here. Other processing steps and data analysis were done by our in-house Python and R scripts.

Data downloaded from TCGA databases were in tab-separated table format and contained processed methylation density at each CpG site. We selected CpG sites that were available in TSMA’s regions and discarded the rest. In summary, we retained around 1088 regions on average. Regions’ values were calculated by the average methylation density values of their CpG sites. With this transformation, our TSMA is technically interchangeable between the sequencing platform and the microarray platform.

### Multi-modal cfDNA feature sets

We adopted a multi-modal set of genome-wide and targeted features from [9] to combine with our deconvolution scores derived from the TSMA. This set included genome-wide methylation density (GWMD), targeted region methylation density (TMD), genome-wide fragmentation profile (GWFP and Flen), end-motif distribution (EM) and copy number aberration (CNA). We refer to [9] for detailed constructions of these features.

### Construction of the methylation reference matrix (atlas)

Reads covering at least one CpG site in a region were collected. Since we were interested in the methylation pattern of CpG sites in the region only, information on other CpG sites carried by the reads was discarded. For a given region *j,* let us denote by 𝑁_𝑗_ the total number of times a CpG site was covered by a read and 𝑀_𝑗_ the number of times a CpG site was covered by a read and was methylated. The “region value” methylation level 𝛽_𝑗_ was then calculated by the ratio of 𝑀_𝑗_ to 𝑁_𝑗_. In regions whose depth of coverage were zero, we discarded the calculation and considered the region as missing input data without applying any imputation technique. For each class of sample and each region, a Student t-test was performed. False discovery rate was controlled by a Bonferroni test correction. Top-500 negative logFC and significantly different regions in each one-versus-rest test were selected. The final tumor-specific methylation matrix is constructed by aggregating the average methylation density of all samples having the same label at each region. Missing values are removed before aggregating. We finally obtained at a methylation reference matrix of shape 6 × 𝑚.

### Non-negative least square matrix factorization

For a given cfDNA sample, to determine the relative weights (deconvolution scores) of each class that contribute to the sample, we implemented a simple non-negative least square (NNLS) algorithm. Let us denote by 𝑋 the methylation reference matrix, 𝑋 ∈ ℝ^6^ ^×𝑚^, representing 6 contributing classes and 𝑚 selected regions. The given input cfDNA sample was represented by a vector 𝛼 of shape 1 × 𝑚. NNLS proceeds to find the weights by solving the following minimization problem.

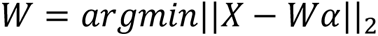

constrained by 𝑊 ≥ 0, where 𝑊 ∈ ℝ^6^ ^×1^. 𝑊 was then normalized to unit sum. To solve this optimization problem, we implemented an in-house Python script based on the Python *scipy* library.

### Graph convolutional neural network (GCNN)

Following the same procedure as our previous work [9], we constructed a graph to train the GCNNs model from discovery and validation cohorts, comprised of low-depth WGBS cfDNA samples of five cancer types (Dataset 4). The overall framework was depicted in [9]. The discovery cohort was then split into two subgroups, including the train dataset and validation dataset for 10-fold cross-validation with stratified sampling to ensure that each cancer class within the train dataset and validation dataset receives the proper representation. The model achieved the highest accuracy among ten folds on the validation dataset chosen and evaluated on an unseen test dataset built from the validation cohort. The undirected input graph 𝐺 = (𝑉, 𝐸) incorporated a node set 𝑉 = {𝑋_𝑖_, 𝑌_𝑖_ |𝑖 = 1, …, 𝑁} (|𝑉| = 𝑁) and an edge set 𝐸 = {𝑒_𝑖𝑗_} (|𝐸| = 𝜀), where 𝑋_𝑖_ and 𝑌_𝑖_ denoted a node 𝑖 and its label, 𝑁 and 𝜀 denoted the number of node and edge in the graph, respectively. Each node 𝑋_𝑖_ was represented by a feature vector 𝑥_𝑖_ ∈ ℝ^𝑑^, which was a concatenation of groups of features (e.g. GWMD, Deconvolution score). We constructed interconnections between nodes at the first layer by k-nearest neighbor (𝑘-NN) of 𝚾 = {𝑥_𝑖_|𝑖 = 1, …, 𝑁} ∈ ℝ^𝑁×𝑑^, where 𝑘 = 5 in our experiments. We defined an adjacency matrix 𝐴 = {𝑎_𝑢𝑣_} ∈ ℝ^𝑁×𝑁^ where we initialized 𝑎_𝑢𝑣_ = 1 if an edge (𝑢, 𝑣) ∈ 𝐸, and 𝑎_𝑢𝑣_ = 0 otherwise. While the label 𝑌_𝑖_ ∈ {𝑡𝑟𝑎𝑖𝑛 𝑑𝑎𝑡𝑎𝑠𝑒𝑡, 𝑣𝑎𝑙𝑖𝑑𝑎𝑡𝑖𝑜𝑛 𝑑𝑎𝑡𝑎𝑠𝑒𝑡} was seen by the model during the training phase, the label 𝑌_𝑗_ ∈ {𝑡𝑒𝑠𝑡 𝑑𝑎𝑡𝑎𝑠𝑒𝑡} was unseen, denoted as unknown nodes, and only used in the evaluation phase. In this way, we adopted a transudative learning [39] setting to analyze the model’s performance on the TOO prediction.

Regarding the model’s architecture, we based our model on Graph Transformer Networks [40], a graph convolutional neural network with support for learning interconnections between nodes in latent spaces. At every layer 𝑙_𝑖_ ∈ {𝑙_1_, …, 𝑙_𝐿_ }, feature representations of node 𝑥_𝑢_ at layer 𝑙_𝑖_ and edges connected node 𝑢 and node 𝑣 were defined by:

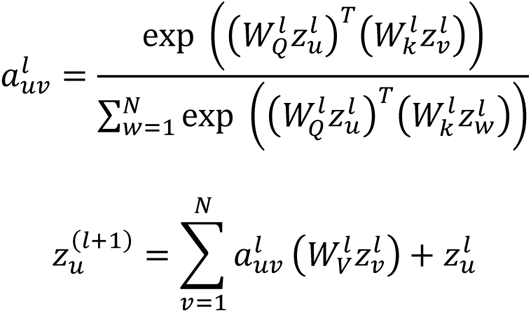

where 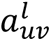 was the connection between node 𝑢 and node 𝑣 at the 𝑙-th layer, 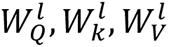 were the learnable parameters at the 𝑙-th layer, 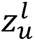 was the latent feature vector of node 𝑢 at the 𝑙-th layer. In this work, we chose 𝐿 = 2, each with the hidden feature size of 64. The output prediction layer was a single Multi Linear Perceptron (MLP).

We dealt with the TOO prediction as a node classification problem, where each node was predicted as one of the five classes: Breast, CRC, Gastric, Liver, and Lung. To avoid the model bias towards major class for the imbalance dataset problem, we applied the focal loss [41] to guide the model focusing more on hard, misclassified samples. The model was trained using the Adam optimizer [42] with a learning rate of 10^−3^. A general description of our model’s architecture is shown in Supplementary Figure 3.

## Results

### Construction of a tumor-specific methylation atlas (TSMA)

The construction of the TSMA relied on an established hypothesis that adjacent CpG sites could be co-methylated and share similar methylation status [18], [25]–[29]. We therefore defined a CpG region as a region of 100bp in length covering at least 5 CpG sites. This definition allowed us to capture regions of dense CpG sites and multiple CpG sites within a region were expected to be covered by a single sequencing read. With this definition, we identified approximately 1.1 million CpG regions in the human reference genome hg19 (Supplementary table 1). We then used WGBS data of 64 tumor tissue samples and 24 WBC samples from the Atlas construction dataset (Figure 1, Materials and methods) to calculate the “region value”, defined as proportion of total methylated CpGs in the reads mapped to a region over the total CpGs that covered by these reads (Figure 1), for each CpG region. To construct the TSMA, the regions in which region values were significantly different among five cancer tissues and WBC were captured. This process allowed the selection of the top 500 regions with the highest absolute log2 fold-change (log2FC) using a one-versus-rest test statistics strategy. Only regions with negative log2FC (significantly lower in the test tissue versus rest) were chosen, based on the observation that most cell-type specific differentially methylated CpG regions were unmethylated [24]. These selected regions were then organized into their respective groups and sorted based on their absolute log2FC values. Representative values for each tissue type at each region was calculated by averaging the region value across all respective samples, as illustrated in Figure 2A. Overall, a TSMA of 2,945 differential CpG regions between 5 tumor tissue types and WBC were constructed.

**Figure 2.**
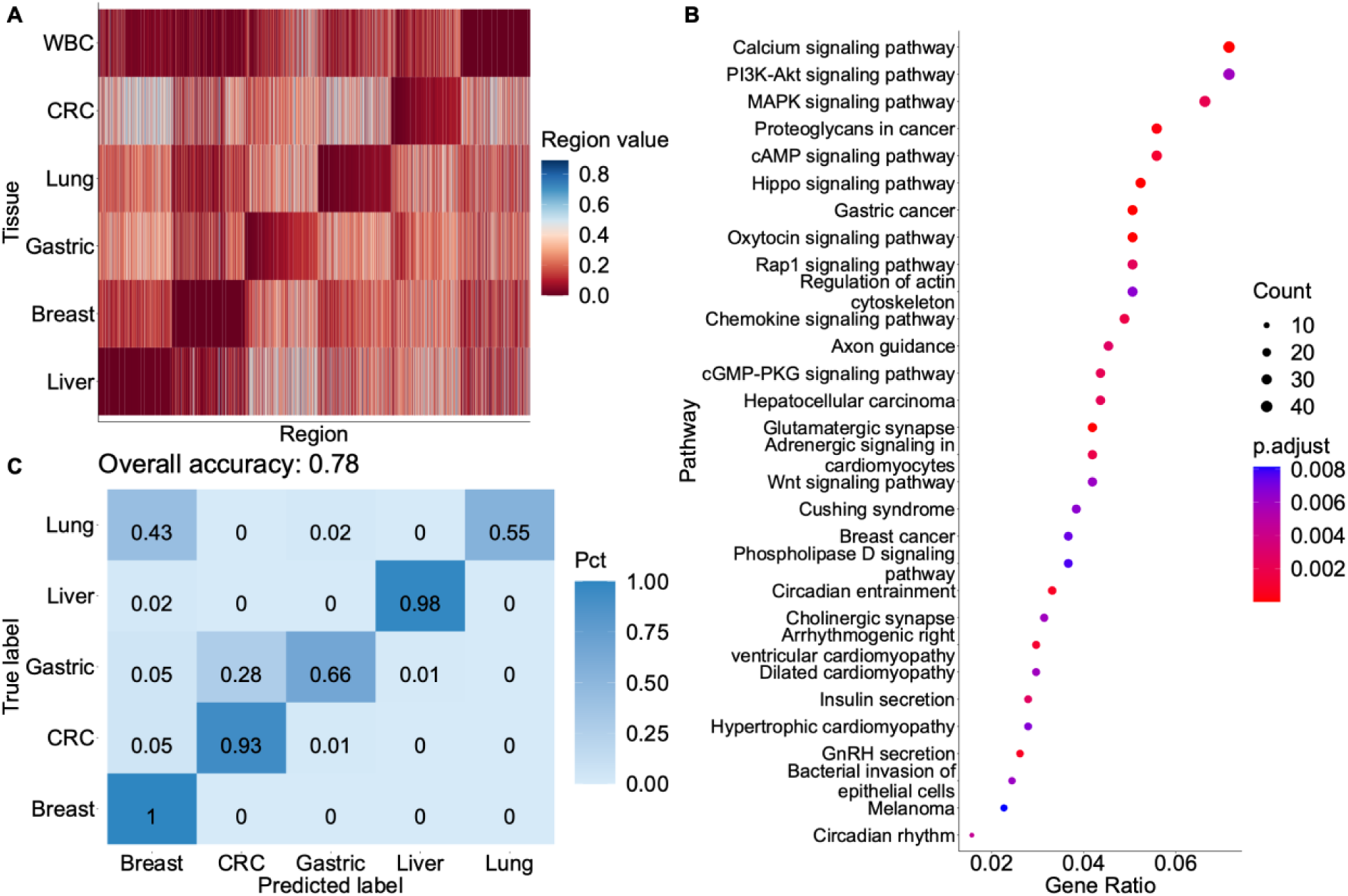
The tumor-specific methylation atlas. (A) Heatmap of average region values in each cancer tissue type or WBC across 2,945 CpG regions included in the TSMA. (B) Pathway analysis reveals cancer-related pathways, which were enriched by the set of genes to which TSMA regions were mapped. (C) Prediction performance using highest deconvolution score to assign label to samples in Dataset 1 comprising of 888 colorectal samples, 1,814 lung samples, 398 gastric samples, 888 breast samples and 429 liver samples.

To assert that our methodology could select for cancer relevant regions, we mapped these regions to genes and conducted a gene set over-representation analysis using GO and KEGG databases to identify pathways that were enriched by our selected regions. Indeed, cancer-related pathways emerged within the top 30 enriched terms (Figure 2B). This result provided the first layer of evidence that our TSMA could capture tumor-specific methylation signals. Encouraged by these outcomes, we next hypothesize that our TSMA could be used to deconvolute samples from an independent source with methods of measuring methylation signal not limited to WGBS. To test this hypothesis, we obtained the 450K/850K methylation microarray data comprising of 4,415 samples of cancer tumor tissues from the 5 cancer types of interest from TCGA database [30] (Dataset 1, **Materials and methods**). We transformed the CpG-wise microarray data into region-wise data to conform with our configuration (Materials and methods). We then performed deconvolution by NNLS and the label of sample was assigned by its highest deconvolution score component. This resulted in an overall accuracy of 78% (Figure 2C). Specifically, we achieved accuracies of 100%, 98% and 93% for breast, liver and CRC cancer, respectively; while gastric and lung cancer exhibited lower accuracies of 66%, and 55%. These results validated our hypothesis and suggested that our TSMA has successfully captured cancer-specific signals that could be used to determine the TOO of a sample.

### Significant correlation between deconvolution scores from TSMA and proportion of tumor DNA

Deconvolution scores derived from a DNA methylation atlas of normal human cell types provided estimates of underlying proportion for each cell type within a sample [31]. In the case of our TSMA, we expected that the TSMA-derived deconvolution scores would estimate the proportion of cancer DNA fragments corresponding to the five specific cancer types used to build TSMA. To confirm this, we first generated a set of samples with known amount of ctDNA by *in silico* mixing DNA fragments (WGBS reads) from tumor tissue into three different cfDNA WGBS samples from healthy donors at various fractions from 0.01% to 25% (Dataset 2, Materials and methods). We then evaluated the correlation between deconvolution scores and the known abundances of tissue-derived DNA fragments. In all samples, the proportions of WBC were consistently reported as the majority and decreased as the amount of tumor tissue DNA increased (Supplementary Figure 1).

We observed that at extremely low tumor fraction (≤ 0.1%) deconvolution scores were mostly 0, except for liver and gastric tumor where the scores were higher but not changing at 0.01%, 0.05% and 0.1%, suggesting a limit for this method at ∼0.1% (Figure 3). At tumor fraction higher than 0.1%, deconvolution scores showed good correlation with known tumor fractions (𝑅 > 0.78) (Figure 3).

**Figure 3.**
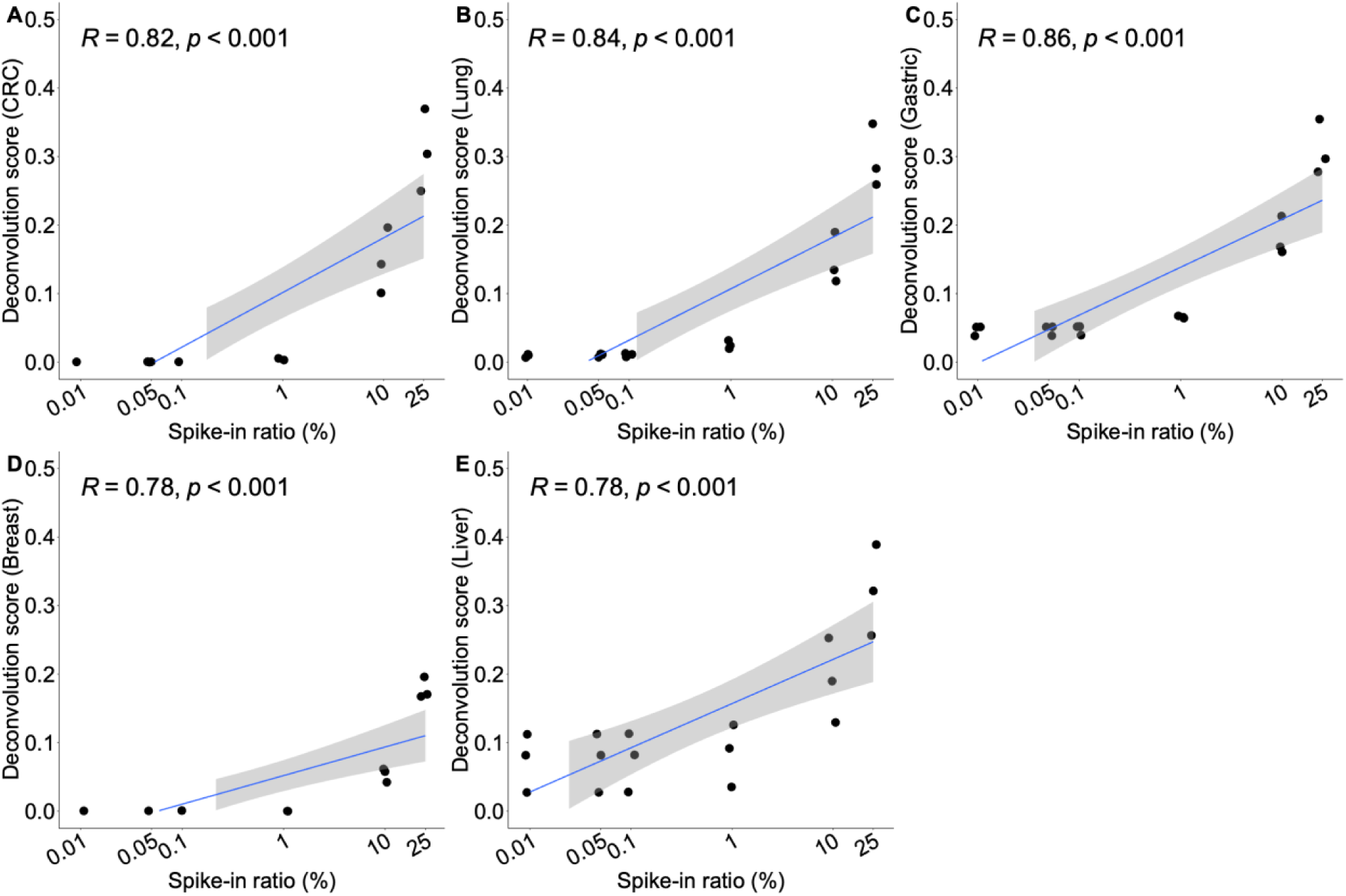
Correlation between deconvolution scores and the proportion of cancer tissue fragments in *in vitro* spike-in samples. Significant correlation (Pearson correlation 𝑅 ≥ 0.78) between spike-in ratios (ie. percent of tumor DNA), ranging from 0.01% to 25%, and deconvolution scores (ie. fractions of specific cell type) was observed in (A) CRC, (B) Lung, (C) Gastric, (D) Breast, and (E) Liver cancer. Spike-in samples were generated by randomly sampling tumor tissue DNA fragments (WGBS reads) and mixing with 3 healthy cfDNA samples at defined ratios (3 replicates at each ratio). Correlation was measured by Pearson coefficient 𝑅. Values on x-axis were log-scaled for visualization purpose only.

The next set of samples that we used to validate the correlation between deconvolution scores and tumor fractions were wet-lab spike-in samples where genomic DNA from each tumor tissue type were mixed with cfDNA from 2 healthy donors at four ratios (0.1%, 1%, 10%, and 25%), the resulting mixed DNA samples were then subjected to WGBS before calculating deconvolution scores (Dataset 3, Materials and methods). Deconvolution scores from our wet-lab spike-in experiment showed similar results with our *in silico* experiment. Across the dataset, we observed correlations in most cancer tissue types with the lowest correlation coefficient (𝑅 = 0.58) in CRC and highest correlation coefficient (𝑅 = 0.87) in Lung cancer (Figure 4). In summary, both *in silico* and wet-lab spike-in experiments demonstrated the ability of TSMA to deconvolute tumor tissue fragments with high correlation to their abundance, suggesting the potential of this approach in predicting TOO.

**Figure 4.**
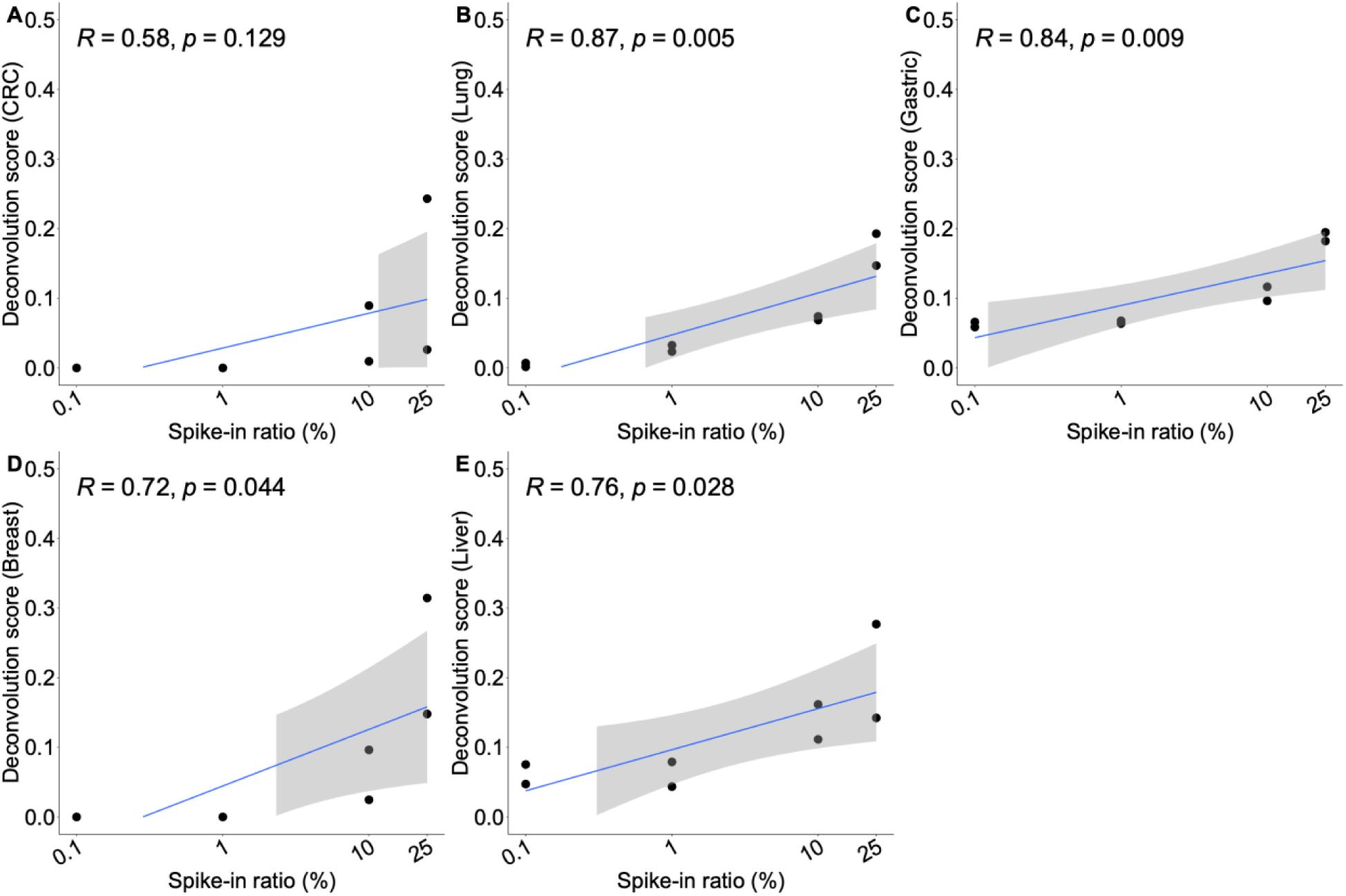
Correlation between deconvolution scores and the proportion of cancer tissue fragments in wet-lab spike-in samples. Strong correlation between spike-in ratios and deconvolution scores was observed in (A) CRC, (B) Lung, (C) Gastric, (D) Breast, and (E) Liver cancer. Wet-lab spike-in samples were generated by mixing genomic DNA from each tumor tissue type with cfDNA from 2 healthy donors at four ratios (0.1%, 1%, 10%, and 25%), before WGBS and calculating deconvolution scores (2 replicates at each ratio). Correlation was measured by Pearson coefficient 𝑅. Values on x-axis were log-scaled for visualization purpose only.

Since changes in deconvolution scores were significantly correlated to tumor abundance in cfDNA sample, we further explored the possibility of using deconvolution scores to determine TOO in a given cfDNA sample [31], [32]. Deconvolution scores of all cfDNA samples in Dataset 2 and 3 consistently showed WBC fractions at ∼90%, leaving only ∼10% for the other five tissue types (Supplementary Figure 1). Therefore, we removed the WBC fraction and used the top tissue type (out of the 5 remaining types) as the TOO classification label. In Dataset 2, we correctly identified the TOO in 119/270 samples (overall accuracy of 44%, Supplementary table 3). However, this performance was strongly affected by the spike-in ratio. The accuracy rose to 92% (Supplementary Figure 2A) for samples with spike-in ratios greater than or equal to 10% (83/90 samples were correctly identified), and dropped to only 20% (Supplementary Figure 2B) for samples with spike-in ratios less than 10% (36/180 correctly identified). Similarly, TOO prediction was correct in 22/40 samples of Dataset 3 (overall accuracy of 55%, Supplementary table 3), of which the samples with spike-in ratios greater than or equal to 10% exhibited 75% accuracy (Supplementary Figure 2C), compared to accuracy of 35% for samples with spike-in ratios less than 10% (Supplementary Figure 2D). Thus, our data indicated that the deconvolution approach worked well only in samples where the tumor tissue abundance exceeded 10%.

### Combining deconvolution scores and genome-wide methylation density in a graph convolutional neural network enhances prediction performance

In early cancer and TOO detection from cfDNA, the multi-modal approach has become increasingly popular, where different features derived from different characteristics of cfDNA are combined as inputs for a machine learning or deep learning model to improve performance [7]– [12]. We have previously published a GCNN model using methylomics, fragmentomics, copy number, and end motifs features for TOO detection with encouraging accuracy of 70% [9]. Here, we used the same dataset (Dataset 5) [9] to explore the possibility of combining deconvolution scores with other cfDNA features to enhance TOO performance. In total, we examined 57 combinations of deconvolution scores (Supplementary table 4) with genome-wide methylation density (GWMD), targeted methylation density (TMD), copy number aberration (CNA), and end motifs (EM) to search for the best performing model. For each combination, we trained a GCNN in a set of 438 cancer patient samples and validated the model in a held-out 239 samples (Dataset 5). GWMD, expressed as the average methylation density across non-overlapping 1M bins in the entire genome, when combined with deconvolution scores achieved the highest accuracy of 69% (Figure 5A, D). This was markerly improved from deconvolution scores alone (26% accuracy, Figure 5B), or GWMD alone (63% accuracy, Figure 5C). This result highlighted the contribution of TSMA deconvolution scores in the GCNN model for TOO detection, especially when combined with GWMD.

**Figure 5.**
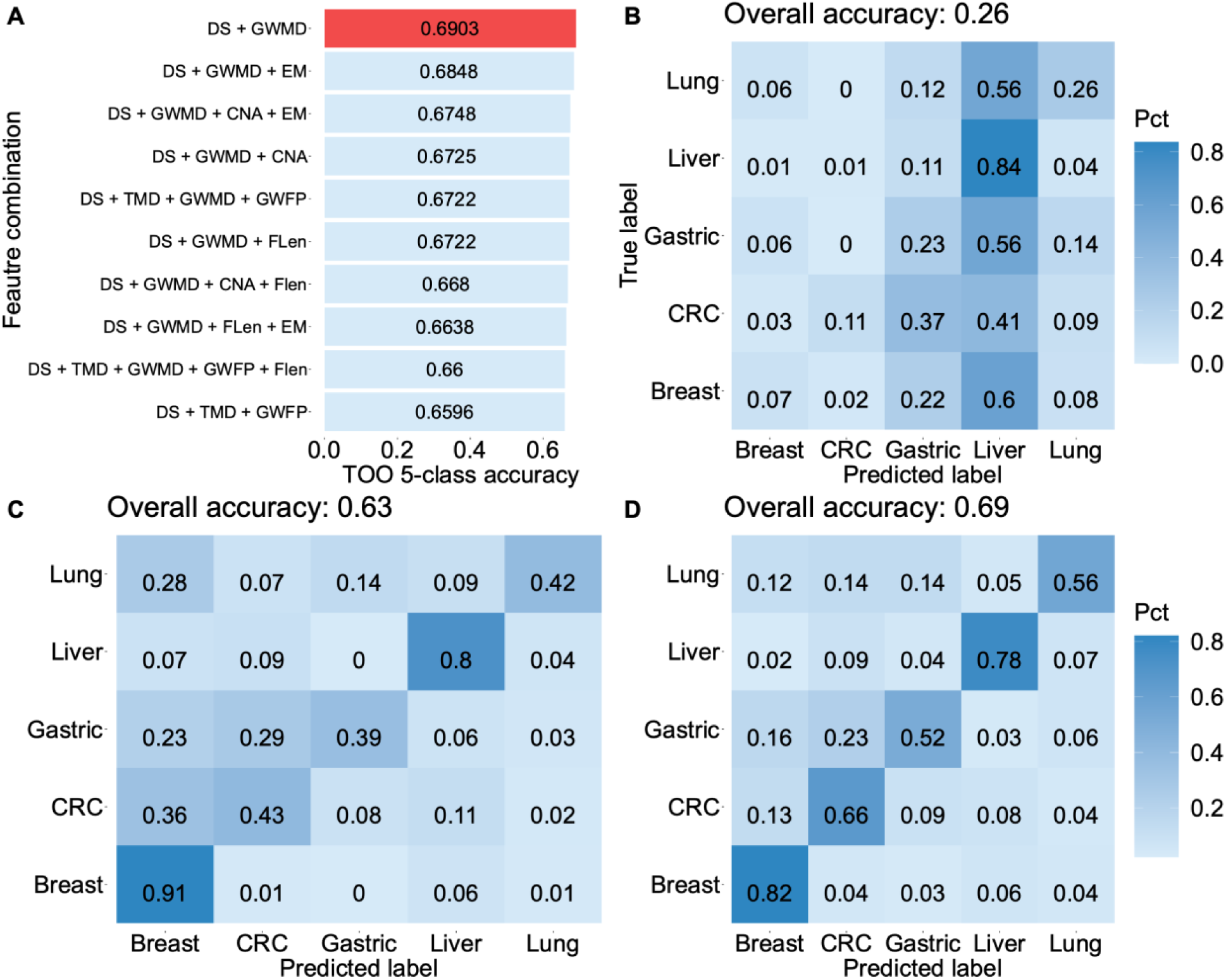
Multi-modal approach combining TSMA deconvolution scores with other cfDNA features in a graph convolutional neural network. **(A)**. Top 10 feature combinations achieving the highest accuracies. Results from other combinations are given in the Supplementary Table 4. The combination of deconvolution score (DS) and GWMD achieved highest accuracy and is shown in red. (B). Confusion matrix obtained from the deconvolution scores. A sample label was assigned by the highest tumor tissue type as indicated by deconvolution scores. (C) Confusion matrix obtained from a GCNN using GWMD feature only. (D) Confusion matrix obtained from a GCNN using both deconvolution scores and GWMD feature.

## Discussion

The multi-modal approach has become increasingly popular for the construction of a machine learning or deep learning model in early cancer and TOO detection from cfDNA [7]–[12]. This approach relies on bioinformatic analysis and feature engineering of WGS or WGBS data to reveal different characteristics of cfDNA, including but not limited to methylomics, fragmentomics, copy number, and end motifs [9]. In this study, we aimed to engineer a new feature that can be used either alone or more likely in combination with other features to enhance the performance of TOO detect from shallow WGBS of cfDNA samples. We constructed the TSMA to distinguish five tumor tissue types and white blood cells using WGBS of tumor tissue and paired WBC samples (Figure 1). This novel approach set our atlas apart from the methylation atlas previously built using healthy human cell types [23], [24]. With this TSMA, methylation data (either WGBS or methylation microarray) can be deconvoluted by non-negative least square matrix factorization into deconvolution scores, which represent the proportion of each tissue type in a sample (Figure 1). Given an input sample, deconvolution scores could be directly utilized to predict its cell type composition, limited to the five tumor tissues and WBC components in our atlas, or used in combination of other features in a model for TOO detection.

In our *in silico* and wet-lab spike-in experiments, we observed strong correlation between deconvolution scores and the known percentage of spike-in tumor-tissue DNA fragments in cfDNA samples (Figure 3 and Figure 4). However, the applications of deconvolution scores in real cfDNA samples, especially in low-depth WGBS samples, posed a greater challenge. In most samples, we found that WBC accounted for nearly 90% of the composition, leaving the sum of all potential tumor-specific signals to less than 10%. This finding aligned with the biological notion that tumor-derived DNA fraction could account for at most 1-10% in cfDNA context [33]. Validations on *in silico* and wet-lab spike-in datasets (Dataset 2, 3) indicated that the limit of detection was 10% tumor abundance to accurately predict a sample TOO (accuracy of 75% to 92%, Figure 3 and Figure 4), which made deconvolution scores alone unsuitable for TOO detection for low-depth cfDNA samples.

Alternatively, deconvolution scores can be used in combination with other cfDNA features readily available from previous works [7]–[9]. Using the same dataset that was presented previously [9], we calculated the deconvolution scores using our newly built TSMA and concatenated them to 57 different combinations of feature vectors (Figure 5A). This combined feature vector was fed to a graph convolutional neural network to achieve the final prediction of the tissue of origin. Training was done on a cohort of 498 low-depth WGBS cfDNA samples (0.5x) and validation on a cohort of 239 low-depth WGBS cfDNA samples (0.5x) of cancer patients. We achieved a 5-class accuracy of 69% for TOO prediction, which is comparable to the results obtained in our previous study [9]. Specifically, compared to the GCNN previously published [9], this new GCNN model achieved higher accuracy for breast and liver (82% vs 78%, and 78% vs 76%), similar accuracy for CRC (66%) and lower accuracy for gastric and lung (52% vs 55%, and 56% vs 63%). This result highlighted that the GCNN using deconvolution scores and GWDM achieved comparable performance to a GCNN built with 9 different sets of features, suggesting that the contribution of deconvolution scores is equal to 8 other feature sets.

There are two main limitations in this work. First, the TSMA was constrained by a dataset of five tumor tissue types and a single set of 24 WBC samples. Future work with more tissue types is needed to expand this atlas and provide a more comprehensive TOO detection. Second, despite achieving an overall accuracy of 69%, there is a noticeably imbalance in the accuracies across different classes of cancer. The unequal distribution of samples across our five cancer types may have introduced bias into our training and validation framework [9]. To further enhance the performance of our approach and mitigate the imbalance, an increase in the number of samples for the underrepresented cancer type is required.

## Conclusions

In conclusion, we have developed a TSMA depicting differential methylated regions across five cancer tumor types and white blood cells. The deconvolution scores from our atlas correlated well with the tumor fraction in cfDNA samples although this is limited mostly to tumor fraction of more than 10%. However, the combination of the deconvolution scores and genome-wide methylation density features significantly enhanced the TOO detection performance when applying to low-depth WGBS cfDNA samples. In summary, our study has paved the way for the application of tumor-specific atlas in TOO detection. Future development of such an atlas might hold the key to significant improvement in TOO detection of all cancer types in low-depth cfDNA samples.

## List of abbreviations

cfDNA: Cell free DNA
TOO: Tissue-of-origin
TSMA: tumor-specific methylation atlas
WBC: white blood cells
NNLS: non-negative least square matrix factorization
GCNN: graph convolutional neural network
GWMD: genome-wide methylation density
TMD: targeted region methylation density
GWFP: genome-wide fragmentation profile
EM: end-motif
CNA: copy number aberration

## Declarations

### Ethics approval and consent to participate

This study was approved by the Ethics Committee of the Medic Medical Center, University of Medicine and Pharmacy and Medical Genetics Institute, Ho Chi Minh city, Vietnam. Written informed consent was obtained from each participant in accordance with the Declaration of Helsinki.

### Consent for publication

Not applicable.

### Availability of data and materials

Analytic data are available on request to the corresponding authors (THN, LST, MDP). Raw FASTQ data are not publicly available due to ethical and regulatory restrictions.

### Competing interests

THN, NHN, HG, LST, MDP receive compensation and have an equity interest in Gene Solutions. NNTD, THT, VTCN, THHN, GTHN are employees of Gene Solutions. The authors ensure that this does not alter the accuracy or integrity of the manuscript. The study was funded by Gene Solutions. The sponsor has no role in the analysis of the data and the preparation of the manuscript.

### Funding

The study was funded by Gene Solutions

### Disclosure statement

We confirm that this does not alter our adherence to Cancer Investigation policies on sharing data and materials.

### Author contribution

Conceptualization: Trong Hieu Nguyen, Hoa Giang, Le Son Tran, Minh Duy Phan

Formal analysis: Nhu Nhat Tan Doan, Trung Hieu Tran, Van Thien Chi Nguyen

Data analysis: Thi Hue Hanh Nguyen

Funding acquisition: Hoai-Nghia Nguyen

Writing-original draft: Trong Hieu Nguyen

Writing-review and editing: Trong Hieu Nguyen, Giang Thi Huong Nguyen, Hoa Giang, Le Son

Tran, Minh Duy Phan

---

**Supplementary Figure 1.**
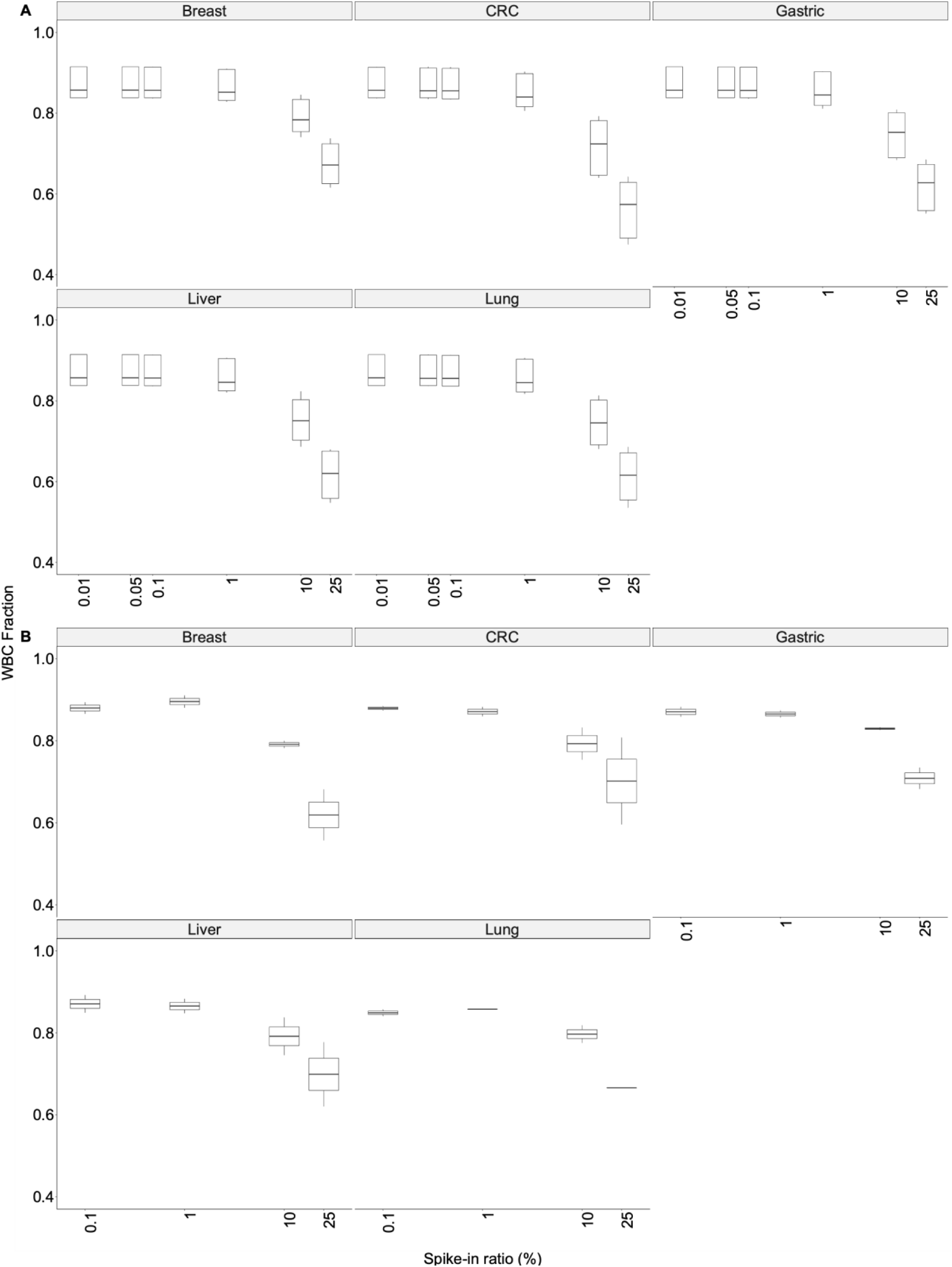
White-blood-cell (WBC) fractions estimated by deconvolution scores and the TSMA in two experimental validation datasets. WBC fractions and tumor spike-in ratios demonstrated an inverse correlation. (A). Inverse correlation between WBC fractions and spike-in ratios in in silico spike-in dataset (Dataset 2). (B). Inverse correlation between WBC fractions and spike-in ratios in wet-lab spike-in dataset (Dataset 3).

**Supplementary Figure 2.**
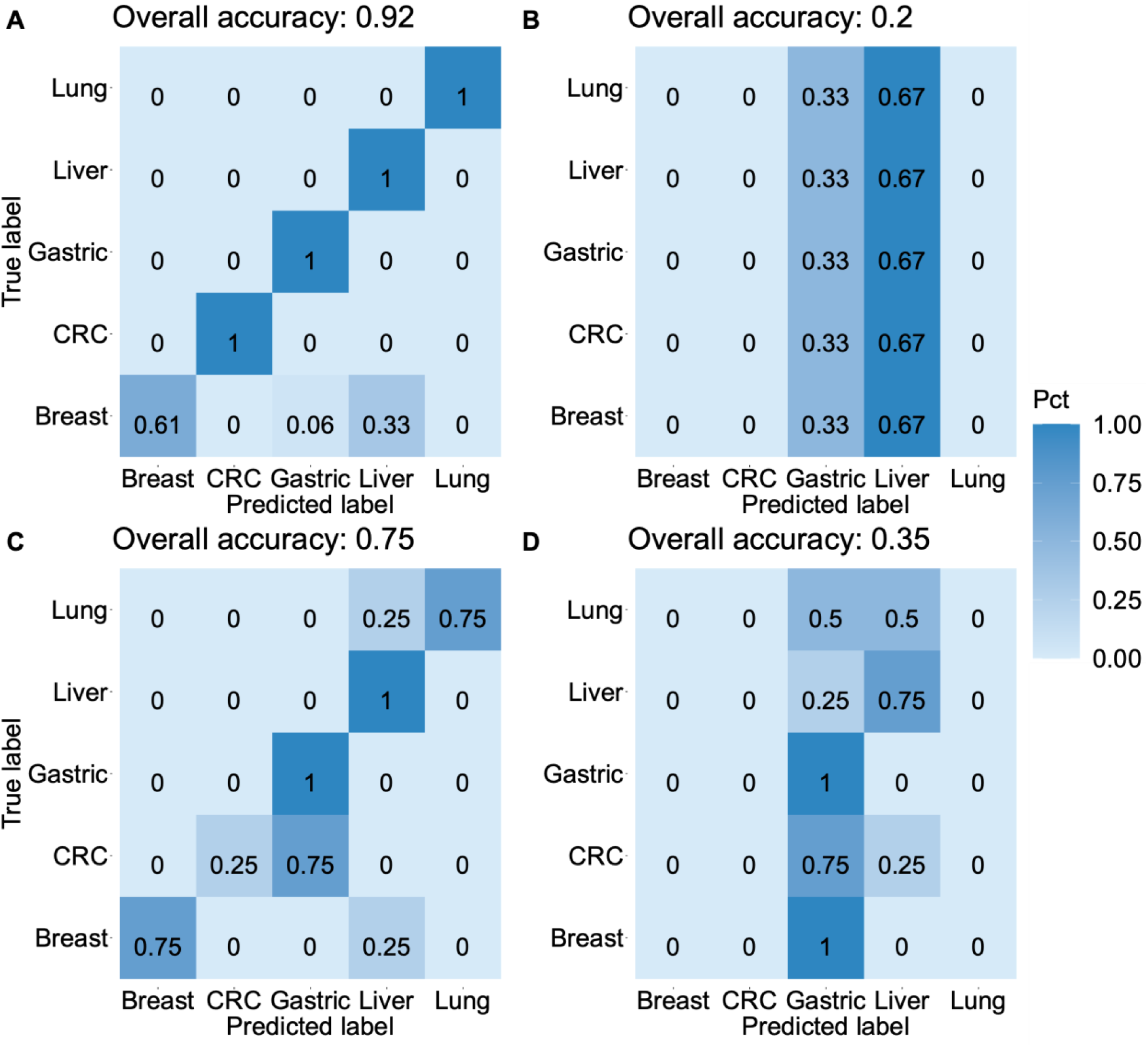
Evaluation of the deconvolution approach in predicting TOO labels of samples in Dataset 2 and 3. Perfomances of the approach were assesed in samples whose spike-in ratio ≥ 10% and spike-in ratio ≤ 10%, separately. (A-B). Confusion matrices obtained from Dataset 2 at spike-in ratio ≥ 10% and spike-in ratio ≤ 10%, respectively. (C-D). Confusion matrices obtained from Dataset 3 at spike-in ratio ≥ 10% and spike-in ratio ≤ 10%, respectively.

**Supplementary Figure 3.**
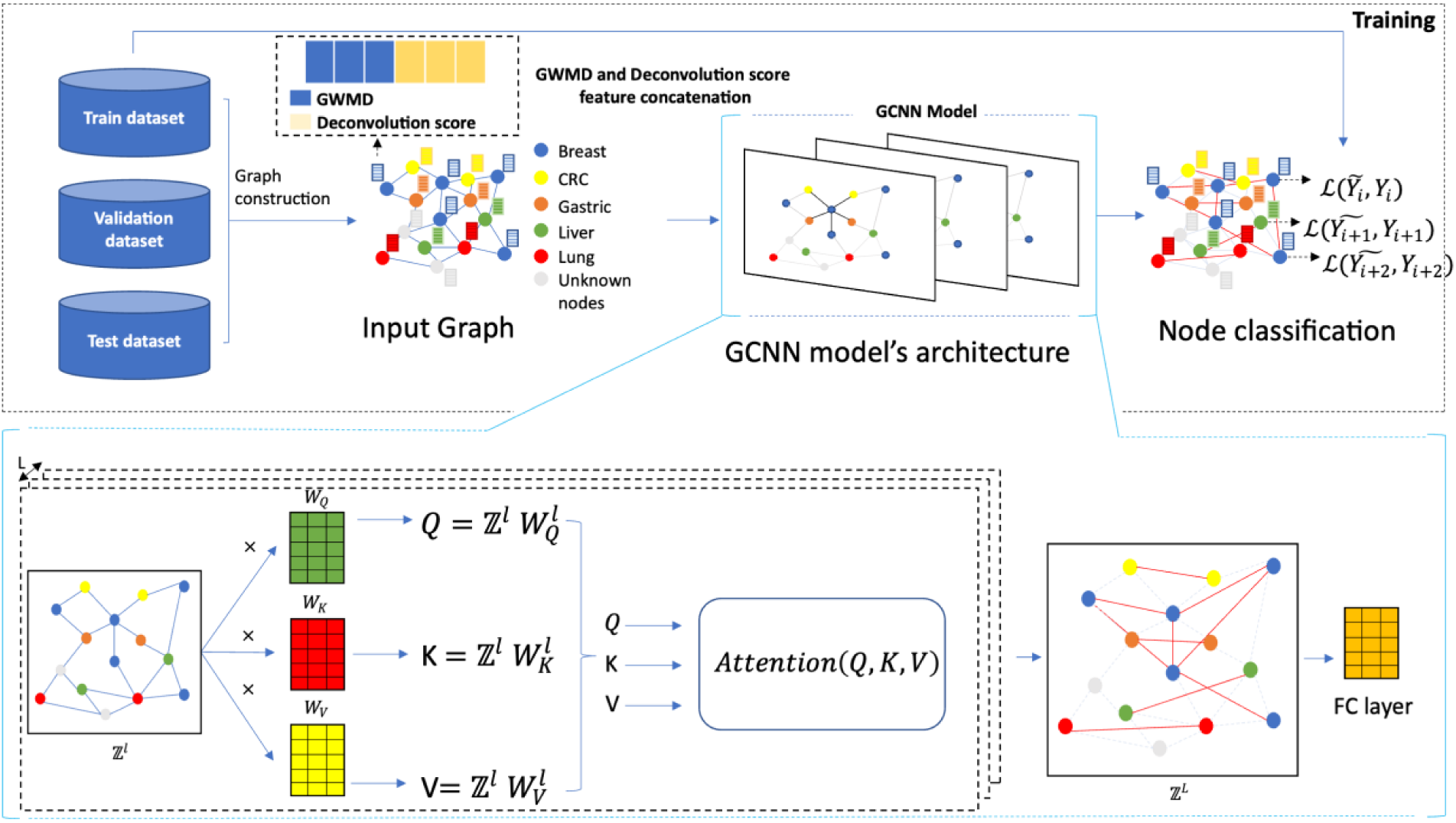
The architect and training scheme of the graph convolutional neural network using deconvolution scores and GWMD feature.

## Supplementary Table

**Supplementary Table 1:** List of 2,945 selected regions in Tumor-specific methylation atlas (TSMA).

**Supplementary Table 2:** Metadata 64 tumor tissue and 24 paired white-blood cell samples.

**Supplementary Table 3:** Metadata of validation datasets (Dataset 1-3) and GCNN model construction and validation dataset (Dataset 4).

**Supplementary Table 4:** Summary of all 57 combinations between deconvolution scores and other cfDNA features.

## References

[1] S. Cristiano et al., “Genome-wide cell-free DNA fragmentation in patients with cancer,” Nature, vol. 570, no. 7761, pp. 385–389, Jun. 2019, doi: 10.1038/s41586-019-1272-6.

[2] V. C. Nguyen et al., “Fragment length profiles of cancer mutations enhance detection of circulating tumor DNA in patients with early-stage hepatocellular carcinoma,” BMC Cancer, vol. 23, no. 1, Dec. 2023, doi: 10.1186/s12885-023-10681-0.

[3] F. Mouliere et al., “Enhanced detection of circulating tumor DNA by fragment size analysis,” 2018. [Online]. Available: https://www.science.org

[4] L. Harbers, F. Agostini, M. Nicos, D. Poddighe, M. Bienko, and N. Crosetto, “Somatic Copy Number Alterations in Human Cancers: An Analysis of Publicly Available Data From The Cancer Genome Atlas,” Front Oncol, vol. 11, Jul. 2021, doi: 10.3389/fonc.2021.700568.

[5] P. Jiang et al., “Plasma DNA end-motif profiling as a fragmentomic marker in cancer, pregnancy, and transplantation,” Cancer Discov, vol. 10, no. 5, pp. 664–673, May 2020, doi: 10.1158/2159-8290.CD-19-0622.

[6] T. H. Phan et al., “Circulating DNA methylation profile improves the accuracy of serum biomarkers for the detection of nonmetastatic hepatocellular carcinoma,” Future Oncology, vol. 18, no. 39, pp. 4399–4413, Dec. 2022, doi: 10.2217/fon-2022-1218.

[7] H. T. Nguyen et al., “Multimodal analysis of ctDNA methylation and fragmentomic profiles enhances detection of nonmetastatic colorectal cancer,” Future Oncology, vol. 18, no. 35, pp. 3895–3912, Dec. 2022, doi: 10.2217/fon-2022-1041.

[8] T. M. Q. Pham et al., “Multimodal analysis of genome-wide methylation, copy number aberrations, and end motif signatures enhances detection of early-stage breast cancer,” Front Oncol, vol. 13, 2023, doi: 10.3389/fonc.2023.1127086.

[9] V. T. C. Nguyen et al., “Multimodal analysis of methylomics and fragmentomics in plasma cell-free DNA for multi-cancer early detection and localization,” Sep. 2023, doi: 10.7554/elife.89083.2.

[10] A. Jamshidi et al., “Evaluation of cell-free DNA approaches for multi-cancer early detection,” Cancer Cell, vol. 40, no. 12, pp. 1537–1549.e12, Dec. 2022, doi: 10.1016/j.ccell.2022.10.022.

[11] S. Y. Kim et al., “Cancer signature ensemble integrating cfDNA methylation, copy number, and fragmentation facilitates multi-cancer early detection,” Exp Mol Med, Nov. 2023, doi: 10.1038/s12276-023-01119-5.

[12] Y. Li et al., “Multi-omics integrated circulating cell-free DNA genomic signatures enhanced the diagnostic performance of early-stage lung cancer and postoperative minimal residual disease,” EBioMedicine, vol. 91, May 2023, doi: 10.1016/j.ebiom.2023.104553.

[13] S. Kang et al., “CancerLocator: Non-invasive cancer diagnosis and tissue-of-origin prediction using methylation profiles of cell-free DNA,” Genome Biol, vol. 18, no. 1, Mar. 2017, doi: 10.1186/s13059-017-1191-5.

[14] W. Li et al., “CancerDetector: ultrasensitive and non-invasive cancer detection at the resolution of individual reads using cell-free DNA methylation sequencing data,” Nucleic Acids Res, vol. 46, no. 15, p. E89, Jun. 2018, doi: 10.1093/NAR/GKY423.

[15] M. C. Liu et al., “Sensitive and specific multi-cancer detection and localization using methylation signatures in cell-free DNA,” Annals of Oncology, vol. 31, no. 6, pp. 745–759, Jun. 2020, doi: 10.1016/j.annonc.2020.02.011.

[16] G. Egger, G. Liang, A. Aparicio, and P. A. Jones, “Epigenetics in human disease and prospects for epigenetic therapy,” Nature, vol. 429, no. 6990, pp. 457–463, 2004, doi: 10.1038/nature02625.

[17] A. P. Feinberg and B. Tycko, “The history of cancer epigenetics,” Nat Rev Cancer, vol. 4, no. 2, pp. 143–153, 2004, doi: 10.1038/nrc1279.

[18] R. Jaenisch and A. Bird, “Epigenetic regulation of gene expression: How the genome integrates intrinsic and environmental signals,” Nature Genetics, vol. 33, no. 3S. pp. 245– 254, 2003. doi: 10.1038/ng1089.

[19] Y. Dor and H. Cedar, “Principles of DNA methylation and their implications for biology and medicine,” The Lancet, vol. 392, no. 10149, pp. 777–786, Sep. 2018, doi: 10.1016/S0140-6736(18)31268-6.

[20] R. Lehmann-Werman et al., “Identification of tissue-specific cell death using methylation patterns of circulating DNA,” Proc Natl Acad Sci U S A, vol. 113, no. 13, pp. E1826–E1834, Mar. 2016, doi: 10.1073/pnas.1519286113.

[21] K. Sun et al., “Plasma DNA tissue mapping by genome-wide methylation sequencing for noninvasive prenatal, cancer, and transplantation assessments,” Proc Natl Acad Sci U S A, vol. 112, no. 40, pp. E5503–E5512, Oct. 2015, doi: 10.1073/pnas.1508736112.

[22] S. Y. Shen et al., “Sensitive tumour detection and classification using plasma cell-free DNA methylomes,” Nature, vol. 563, no. 7732, pp. 579–583, Nov. 2018, doi: 10.1038/s41586-018-0703-0.

[23] J. Moss et al., “Comprehensive human cell-type methylation atlas reveals origins of circulating cell-free DNA in health and disease,” Nat Commun, vol. 9, no. 1, Dec. 2018, doi: 10.1038/s41467-018-07466-6.

[24] N. Loyfer et al., “A DNA methylation atlas of normal human cell types,” Nature, vol. 613, no. 7943, pp. 355–364, Jan. 2023, doi: 10.1038/s41586-022-05580-6.

[25] F. Eckhardt et al., “DNA methylation profiling of human chromosomes 6, 20 and 22,” Nat Genet, vol. 38, no. 12, pp. 1378–1385, Dec. 2006, doi: 10.1038/ng1909.

[26] R. A. Irizarry et al., “Comprehensive high-throughput arrays for relative methylation (CHARM),” Genome Res, vol. 18, no. 5, pp. 780–790, May 2008, doi: 10.1101/gr.7301508.

[27] R. Lister et al., “Human DNA methylomes at base resolution show widespread epigenomic differences,” Nature, vol. 462, no. 7271, pp. 315–322, 2009, doi: 10.1038/nature08514.

[28] R. A. Irizarry et al., “The human colon cancer methylome shows similar hypo- and hypermethylation at conserved tissue-specific CpG island shores,” Nat Genet, vol. 41, no. 2, pp. 178–186, Feb. 2009, doi: 10.1038/ng.298.

[29] S. Guo, D. Diep, N. Plongthongkum, H. L. Fung, K. Zhang, and K. Zhang, “Identification of methylation haplotype blocks AIDS in deconvolution of heterogeneous tissue samples and tumor tissue-of-origin mapping from plasma DNA,” Nat Genet, vol. 49, no. 4, pp. 635– 642, Mar. 2017, doi: 10.1038/ng.3805.

[30] J. N. Weinstein et al., “The cancer genome atlas pan-cancer analysis project,” Nat Genet, vol. 45, no. 10, pp. 1113–1120, Oct. 2013, doi: 10.1038/ng.2764.

[31] A. J. Titus, R. M. Gallimore, L. A. Salas, and B. C. Christensen, “Cell-type deconvolution from DNA methylation: A review of recent applications,” Human Molecular Genetics, vol. 26, no. R2. Oxford University Press, pp. R216–R224, Oct. 01, 2017. doi: 10.1093/hmg/ddx275.

[32] A. Lubotzky et al., “Liquid biopsy reveals collateral tissue damage in cancer,” JCI Insight, vol. 7, no. 2, Jan. 2022, doi: 10.1172/jci.insight.153559.

[33] M. H. D. Neumann, S. Bender, T. Krahn, and T. Schlange, “ctDNA and CTCs in Liquid Biopsy – Current Status and Where We Need to Progress,” Computational and Structural Biotechnology Journal, vol. 16. Elsevier B.V., pp. 190–195, Jan. 01, 2018. doi: 10.1016/j.csbj.2018.05.002.

[34] S. Andrews, “FastQC.” Jun. 2010. [Online]. Available: https://qubeshub.org/resources/fastqc

[35] https://zenodo.org/record/7598955, “TrimGalore”.

[36] F. Krueger and S. R. Andrews, “Bismark: A flexible aligner and methylation caller for Bisulfite-Seq applications,” Bioinformatics, vol. 27, no. 11, pp. 1571–1572, Jun. 2011, doi: 10.1093/bioinformatics/btr167.

[37] “Picard toolkit,” Broad Institute, GitHub repository. Broad Institute, 2018.

[38] P. Danecek et al., “Twelve years of SAMtools and BCFtools,” Gigascience, vol. 10, no. 2, Feb. 2021, doi: 10.1093/gigascience/giab008.

[39] T. Joachims, “Transductive Learning via Spectral Graph Partitioning,” in Proceedings of the Twentieth International Conference on International Conference on Machine Learning, in ICML’03. AAAI Press, 2003, pp. 290–297.

[40] S. Yun, M. Jeong, R. Kim, J. Kang, and H. J. Kim, “Graph Transformer Networks.”

[41] T.-Y. Lin, P. Goyal, R. Girshick, K. He, and P. Dollár, “Focal Loss for Dense Object Detection.”

[42] D. P. Kingma and J. Ba, “Adam: A Method for Stochastic Optimization,” in *3rd International Conference on Learning Representations, ICLR 2015, San Diego, CA, USA, May 7-9*, *2015, Conference Track Proceedings*, Y. Bengio and Y. LeCun, Eds., 2015. [Online]. Available: http://arxiv.org/abs/1412.6980

